# STAT1-dependent tolerance of intestinal viral infection

**DOI:** 10.1101/2020.02.13.936252

**Authors:** Heather A Filyk, Ava J Sharon, Nicolette M Fonseca, Rachel L Simister, Wallace Yuen, Blair K Hardman, Hannah G Robinson, Jung Hee Seo, Joana Rocha-Pereira, Ian Welch, Johan Neyts, Sean A Crowe, Lisa C Osborne

**Author notes:** Authors contributed equally (Heather A Filyk, Ava J Sharon). Lead Contact: Lisa Osborne.

## Abstract

Recent evidence indicates that viral components of the microbiota can contribute to intestinal homeostasis and protection from local inflammatory or infectious insults. However, host-derived mechanisms that maintain tolerance to the virome remain largely unknown. Here, we use colonization with the model commensal murine norovirus (MNV CR6) to interrogate host-directed mechanisms of viral tolerance, and show that STAT1 is a central coordinator of tolerance following CR6 colonization. STAT1 restricts CR6 replication to the intestinal tract, prevents systemic viral-induced tissue damage and disease, and regulates antiviral CD4^+^ and CD8^+^ T cell responses. In contrast to systemic viral pathogens that drive T cell mediated immunopathology in STAT1-deficient mice, our data indicates that loss of CD4^+^ or CD8^+^ T cells and their associated effector functions has no effect on CR6-induced disease. However, therapeutic administration of an antiviral compound to limit viral replication prevented viral-induced tissue damage and death despite ongoing dysregulated antiviral T cell responses. Collectively, our data uncouple the requirement for STAT1-mediated regulation of antiviral T cell responses from innate immune-mediated restriction of viral replication that is necessary for intestinal viral tolerance.

## Introduction

The mammalian intestinal immune system co-evolved with colonizing species, including viruses, archaea, bacteria, fungi, protists and helminths. In the absence of stimuli from all members of this complex community, germ-free (GF) animals exhibit characteristic defects in intestinal vasculature, epithelial morphology, and immune cell development^1^. The mutualistic nature of this relationship is supported by the evolution of both host and microbe-directed tolerance mechanisms that limit the generation of inflammatory responses against the colonizing microbes^2^. Some of the molecular mechanisms the host utilizes to coordinate tolerance toward the vast antigenic diversity encoded by the microbes that make up our multibiome have been identified. For example, in the absence of MyD88, IL-10 or NFIL3 (among others), mice exhibit spontaneous or enhanced susceptibility to intestinal inflammation as a result of impaired tolerance toward bacterial components of the microbiota^3–5^.

In addition to the bacterial microbiome, there is increasing appreciation of the roles other microbial groups play in intestinal homeostasis. Monocolonization of mice with members of the mycobiome or virome can restore some of the defects in germ-free mice, indicating that an intact and diverse community contributes to optimal intestinal function^6, 7^. Broad-spectrum antiviral treatment can enhance DSS-induced colonic inflammation in mice^8^, indicating that even in the presence of a complex microbiota, the virome (composed of bacteriophage, endogenous retroviruses, and eukaryotic viruses) contributes to intestinal homeostasis. Supporting the hypothesis that host-encoded mechanisms regulate and tolerate viral colonization, latent or chronic viral infection can be reactivated by helminth infection in a STAT6 and interferon (IFN)*γ*-dependent manner^9^.

MNV CR6 (CR6) demonstrates features of a mutualistic host-microbe interaction and is an attractive model to interrogate commensal tolerance to a eukaryotic virus. In addition to providing supportive signals in immunosufficient GF mice^7, 10^, this strain can establish persistent colonic infection with continued shedding for at least two months in the absence of any detectable clinical or histopathological signs in wildtype (WT) mice^11^. Once MNV was identified^12^, sentinel screening of animal units across North America and Europe identified its presence in >30% of facilities^13^, raising concerns that its presence or absence could contribute to microbiota-induced differences in disease models. In WT mice, MNV infection has little effect on viral co-infection^14, 15^, but can act as a pathobiont to exacerbate colonic inflammation in genetically susceptible animals^16, 17^. Notably, despite elevated viral loads, no clinical signs of disease have been reported in *Rag1*^-/-^, *Rag2*^-/-^, *Ifnar*^-/-^ and *Ifnlr1*^-/-^ mice colonized with CR6^18–21^, indicating that B & T cells, type I and type III interferons are dispensable for homeostatic tolerance to persistent intestinal viral colonization.

Signal transducer and activator of transcription 1 (STAT1) signaling is critical for immune-mediated resistance to multiple viral pathogens. In weanling mice infected with rotavirus (RV), an IL-22/STAT1-dependent mechanism limits intestinal epithelial cell (IEC)-intrinsic viral replication and RV-induced intestinal pathology^22^. Following vaccinia virus infection, STAT1 supports optimal antiviral CD8^+^ T cell expansion and survival^23, 24^. In contrast, STAT1-deficient mice exhibit increased numbers of both CD4^+^ and CD8^+^ T cells in response to infection with LCMV Armstrong, although these cells do not constrain viral replication^25^. Instead, *Stat1*^-/-^ mice succumb rapidly to CD4^+^ T cell mediated immunopathology^25^. Indeed, tissue damage caused by dysregulated lymphocyte subsets has been reported in the context of multiple viral pathogens, including LCMV Armstrong, influenza and respiratory syncytial virus in mice with impaired STAT1 or type I interferon a receptor (IFNAR) signaling^25–29^. Following LCMV clone 13 infection, STAT1-dependent IFNAR signaling promotes dendritic cell expression of IL-10 and PD-L1 to limit CD4^+^ effector function^30, 31^. These data indicate that STAT1 signaling can act on multiple cell types and effect context-dependent outcomes following viral infection. STAT1-deficient mice infected with an acute, systemic strain of murine norovirus (MNV CW3) exhibit rapid mortality^32^ and independent groups have reported viral-induced disease in *Stat1*^-/-^ mice following infection with persistent strains MNV-O7 and MNV4^33–35^, although the mechanisms that drive MNV-induced disease remain undefined.

Here, we examine the role for STAT1 in maintaining host tolerance to persistent CR6 infection. In our hands, CR6-infected STAT1-deficient mice display weight loss and mortality over a course of 30 days, as opposed to the rapid (<1 week) weight loss and mortality associated with CW3 infection. At 7-10 days post-infection (pi), approximately 25% of STAT1-deficient animals display virus-induced weight loss which correlates with splenomegaly, spleen and liver histopathology, as well as dysregulation of colon-associated microbial communities. CR6 infection also elicits neutrophilia, a Th17-skewed CD4^+^ T cell response and hyperaccumulation of antigen-specific CD8^+^ T cells. These data suggested that STAT1 may maintain enteric viral tolerance by limiting microbial translocation and/or immune-mediated pathology. However, microbiome ablation or attenuation of adaptive immune responses was not sufficient to rescue mice from viral-induced disease. In contrast, therapeutic administration of a viral polymerase inhibitor, 2’-C-methylcytidine (2CMC), limited viral replication and prevented viral-induced disease in STAT1-deficient mice. Notably, neutrophilia, Th17 cell differentiation and accumulation of antigen-specific CD8^+^ T cells was still apparent in CR6-infected *Stat1*^-/-^ mice treated with 2CMC. Thus, in contrast to STAT1’s role in limiting immune-mediated damage to host tissue seen in the context of other viral pathogens, STAT1 directly limits CR6 viral replication. These data effectively uncouple the effects of a dysregulated immune response from virus-induced damage to the host and demonstrate that STAT1 contributes to tolerance of a commensal-like enteric viral infection.

## Results

### STAT1 limits MNV CR6 dissemination and protects against viral-induced disease

To limit the impact of environmental influences on our analysis of STAT1 function in the context of CR6 exposure, we bred *Stat1*^+/-^ x *Stat1*^+/-^ to generate littermate *Stat1^+/+^*, *Stat1^+/-^* and *Stat1^-/-^* (*Stat1*^WT^, *Stat1*^Het^ and *Stat1*^KO^) mice. We took advantage of the clear requirement for STAT1 in response to CW3 infection to test whether *Stat1*^Het^ mice displayed any signs of haploinsufficiency. Consistent with previous observations^12^, *Stat1*^KO^ mice demonstrated rapid weight loss and 100% mortality by day 5 post-CW3 infection (Figure S1A, B). In contrast, 100% of *Stat1*^WT^ and *Stat1*^Het^ mice survived through day 8 post-CW3 infection with no signs of weight loss, mortality or other clinical symptoms (Figure S1A, B). In addition, there was no difference in viral clearance, frequency or number of MNV-specific CD8^+^ T cells between *Stat1*^WT^ and *Stat1*^Het^ littermates (Figure S1C-F), indicating that a single copy of the *Stat1* gene is sufficient for antiviral immunity in the context of MNV CW3 infection. Consistent with this, *Stat1*^WT^ and *Stat1*^Het^ bone marrow derived dendritic cells (BMDCs) expressed similar levels of *Stat1* mRNA (Figure S1G) and both responded to MNV infection by up-regulating gene and protein expression of STAT1 (Figure S1G-I). We thus modified our breeding strategy, using *Stat1*^Het^ x *Stat1*^KO^ mice to generate STAT1-sufficient (*Stat1*^Het^) and STAT1-deficient (*Stat1*^KO^) littermates for the remaining experiments.

Using this breeding strategy, we assessed the role of STAT1 in maintaining tolerance to a commensal-like persistent intestinal virus. Consistent with previous reports of CR6 infection in WT mice ^11^, *Stat1*^Het^ mice had no detectable clinical symptoms through 30 days of CR6 exposure despite a high colonic viral burden (Figure 1A-C), indicating infectious tolerance^2^. In contrast, 75% of *Stat1*^KO^ mice demonstrated clinical signs, including >10% loss of body weight and piloerection; ultimately, 70% succumbed to virus-induced disease (Figure 1A, B). At the time of euthanasia (day 30 pi for *Stat1*^Het^), viral loads in the colon and spleen were measured. In contrast to *Stat1*^Het^ mice, CR6 was readily detectable in splenic tissue of *Stat1*^KO^ mice (Figure 1C), suggesting that STAT1 plays a critical role in maintaining the intestinal restriction of CR6 infection. Moreover, tissue and systemic viral burden correlated with virus-induced mortality in *Stat1*^KO^ mice (Figure 1C). Notably, a fraction of *Stat1*^KO^ mice survived the infection without losing >10% body weight or displaying other signs of virus-induced disease. At 30 days post-CR6 exposure, viral burdens were below the limit of detection in both colon and spleen of these mice (Figure 1C).

**Figure 1:**
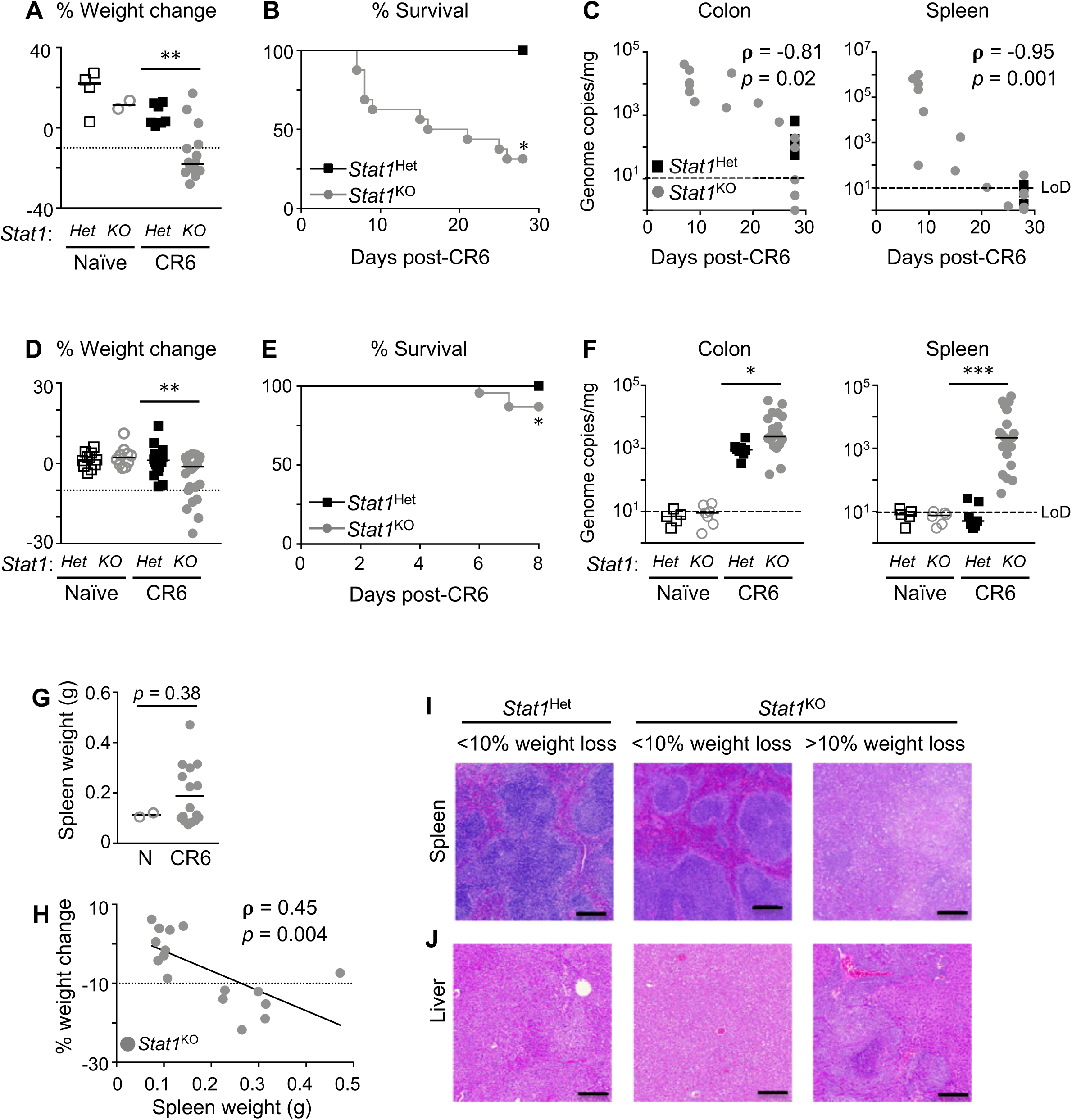
STAT1 limits MNV CR6 dissemination and protects against viral-induced disease. *Stat1^het^* and *Stat1^KO^* mice received 10^4^ pfu MNV CR6 or PBS po and were euthanized on day 28 pi (A-C), day 8 pi (D-F), or at humane endpoint. (A) Weight change as a percentage difference from initial, at day 28 pi or at humane endpoint. (B) Survival curve CR6-infected mice, Mantel-Cox Test. (C) Correlation between MNV CR6 genome copies in colon (left) and spleen (right) and time to humane endpoint in *Stat1^het^* and *Stat1^KO^* mice, Spearman Correlation Coefficient Test. Data in (A-C) are compiled from two independent experiments, n=4 naïve *Stat1^het^*, n=2 naïve *Stat1^KO^*, n=8 *Stat1^het^*, and n=16 *Stat1^KO^* mice. (D) Weight change as a percentage difference from initial, at day 8 pi or at humane endpoint. (E) Survival curve of 8 day CR6-infected mice, Mantel-Cox Test. (F) MNV CR6 genome copies in colon (left) and spleen (right) were quantified by RT-PCR. Data in (D-F) are compiled from six independent experiments, n=11 naïve *Stat1^het^*, n=10 naïve *Stat1^KO^*, n=22 *Stat1^het^*, and n=23 *Stat1^KO^* mice. (G,H) *Stat1^KO^* spleen weight at day 8 pi or humane endpoint (G) and comparison with weight change as percentage difference from initial, Spearman Correlation Coefficient Test (H). (I, J) Representative H&E stained spleen and liver sections at day 8 pi, scale bar = 200 µM. Images are representative of three independent experiments. Two-tailed Mann-Whitney U test comparing *Stat1*^Het^ and *Stat1*^KO^ mice was performed unless otherwise noted. \**P*<0.05, \*\**P*<0.01, ***P<0.001, \*\*\*\**P*<0.0001. LoD: Limit of Detection.

In order to examine mechanisms that could contribute to development of viral-induced disease in these mice, we focused on day 8 post-CR6 exposure, a time point where the majority of *Stat1*^KO^ mice survived the infection, including a portion (∼25%) that had lost >10% of their original body weight (Figure 1D, E). At this time point, *Stat1*^KO^ mice had higher colonic viral burdens than their littermate controls, and viral dissemination to the liver and spleen that correlated with weight loss (Figure 1F, Figure S2A). In *Stat1*^KO^ mice that demonstrated >10% weight loss, there was evidence of virus-induced disease characterized by splenomegaly (Figure 1G, H) and generalized acute multifocal severe necrosis in the spleen and liver that was not evident in CR6-infected healthy *Stat1*^KO^ or *Stat1*^Het^ littermates or in WT mice (Figure 1I, J). Notably, colonic architecture appeared similar between naïve and CR6-infected *Stat1*^KO^ mice (Figure S2B), indicating that infection-induced disease is related to a loss of control of systemic viral replication. Collectively, these data indicate that STAT1-dependent mechanisms restrict systemic viral spread and maintain a tolerance to a commensal-like intestinal virus.

### Distinct patterns of CD4^+^ and CD8^+^ T cell dysfunction in the absence of STAT1

Previous studies indicate that STAT1 can either promote or limit antiviral CD8^+^ T cell accumulation and function^24, 36, 37^. In WT mice, CR6 establishes a replicative niche within intestinal tuft cells where contact with antigen-specific CD8^+^ T cells is limited^18, 38^. However, our data is consistent with other findings that STAT1 and upstream type I IFN signaling are essential to restrict CR6 to this immunoprivileged niche^20^, and we hypothesized that viral dissemination may be caused by an impairment of CD8^+^ T cell function due to the loss of STAT1. Thus, we tested whether the inability of *Stat1*^KO^ mice to restrict viral dissemination correlated with impairments in cytotoxic CD8^+^ T cells. At day 8 post-CR6 infection, we quantified and phenotyped MNV-specific CD8^+^ T cells isolated from the colonic intraepithelial lymphocyte (IEL) and lamina propria lymphocyte (LPL) compartments and the spleen of *Stat1*^Het^ and *Stat1*^KO^ littermates. Similar to WT mice^38^, there was a small but distinguishable population of MNV-specific CD8^+^ T cells in the colonic IEL, LPL and spleen of *Stat1*^Het^ mice (Figure 2A, B). In the spleen, the frequency of MNV-specific CD8^+^ T cells was similar between *Stat1*^Het^ and *Stat1*^KO^ mice, but *Stat1*^KO^ mice demonstrated an increased frequency of colonic IEL and LPL MNV-specific CD8^+^ T cells (Figure 2A, B). Further, stimulation of splenocytes with immunodominant MNV-derived peptides elicited effector molecule (IFN*γ*, TNF*α* and Granzyme B) production from CR6-infected *Stat1*^Het^ and *Stat1*^KO^ mice (Figure 2C), indicating that loss of STAT1 does not impact the accumulation or function of antiviral CD8^+^ T cells in this context. Notably, CD8^+^ T cells isolated from CR6-infected *Stat1^Het^* and *Stat1^KO^* mice both failed to upregulate CD107a and MIP1α, supporting previous studies showing that these cells appear to have suboptimal function in the context of CR6 infection ^38^.

**Figure 2:**
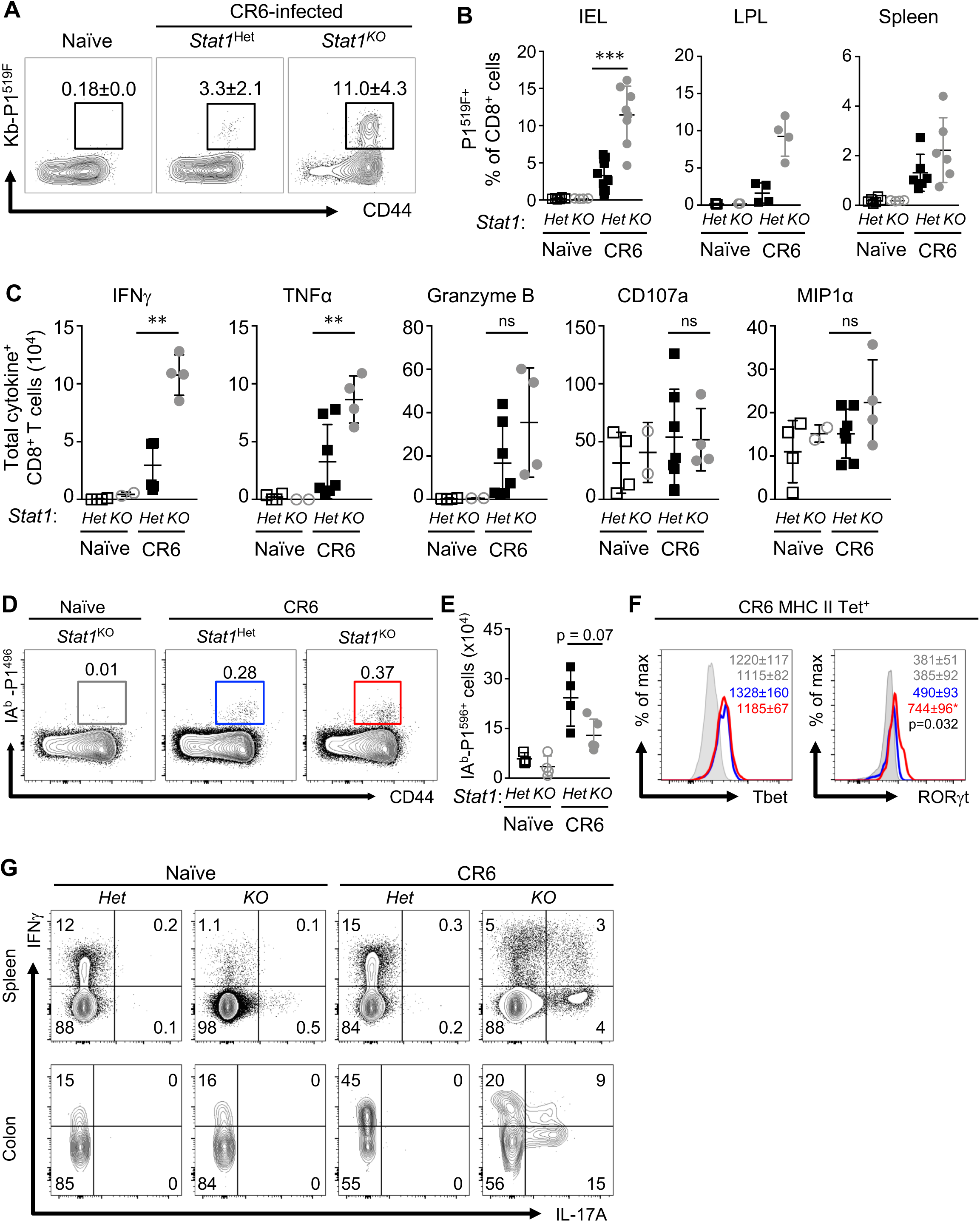
Distinct patterns of CD4^+^ and CD8^+^ T cell dysfunction in the absence of STAT1. *Stat1^het^* and *Stat1^KO^* mice received 10^4^ pfu MNV CR6 or PBS po and were analyzed on day 8 pi. (A) Flow cytometric detection of MNV-CR6 P1^519F^ antigen-specific CD8^+^ T cells isolated from colonic IELs. Numbers in flow plots represent frequencies ± SEM of CR6 Kb-P1^519F^ tetramer^+^ cells within the CD8^+^ lymphocyte gate. (B) Quantification of MNV CR6-specific cells as a frequency of CD45^+^ CD8^+^ IELs, LPLs, and splenocytes. Data in A & B are compiled from three independent experiments, n=4-6 per naïve group and n=12 *Stat1^het^* and n=8 *Stat1^KO^* mice. (C) Total cell counts of IFNγ^+^, TNFα^+^, Granzyme B^+^, CD107a^+^, and MIP1α^+^ CD8^+^ splenocytes following stimulation with MNV P1^519F^ peptide. Data are representative of three independent experiments, n=2-3 per naïve group and n=3-4 per infected group. (D) Lymphocytes from spleen, peripheral and mesenteric lymph nodes naïve and CR6-infected *Stat1*^Het^ and *Stat1*^KO^ mice were pooled and enriched for MHCII IAb-P1^496^ tetramer^+^ cells. Numbers in flow plots represent frequencies ± SEM of MHCII IAb-P1^496^ tetramer^+^ cells within the CD4^+^ lymphocyte gate. (E) Quantification of total IAb-P1^496^ tetramer^+^ CD4^+^ T cells. (F) Representative Tbet and RORγt expression in IAb-P1^496^ tetramer^+^ CD4^+^ T cells. Data in D-F are representative of two independent experiments, n=4 per group. (G) Splenocytes and colonic IELs were stimulated with PMA/ionomycin in the presence of BFA/monensin for 5 hours *ex vivo* and assayed for production of IFN*γ* and IL-17A. Numbers in flow plots represent frequencies ± SEM of cells within the CD4^+^ lymphocyte gate. Data are representative of three independent experiments with n=4-6 mice per group. Two-tailed Mann-Whitney U test comparing *Stat1*^Het^ and *Stat1*^KO^ mice was performed unless otherwise noted. \**P*<0.05, \*\**P*<0.01, ***P<0.001, \*\*\*\**P*<0.0001.

We also characterized the antiviral CD4^+^ T cell response in *Stat1*^Het^ and *Stat1*^KO^ mice via tetramer pulldown of MNV-specific CD4^+^ T cells^39^. In contrast to the increased number of antiviral CD8^+^ T cells detected in *Stat1*^KO^ mice, there was no significant difference in the number of MNV capsid-specific (tetramer IA^b^-P1^496^) CD4^+^ T cells between CR6-infected *Stat1*^Het^ and *Stat1*^KO^ animals (Figure 2D, E). However, MNV-specific *Stat1*^Het^ CD4^+^ T cells expressed T-bet, the master regulator of Th1 differentiation involved in intracellular pathogen immunity, whereas *Stat1*^KO^ CD4^+^ T cells expressed both T-bet and ROR*γ*t (Figure 2F), the transcription factor that specifies Th17 differentiation and is normally associated with protection from extracellular bacterial or fungal pathogens. Consistent with this, polyclonal stimulation of splenocytes or colonic lymphocytes resulted in expression of IFN*γ* in *Stat1*^Het^ mice and robust expression of both IFN*γ* and IL-17A in *Stat1*^KO^ mice (Figure 2G). These data suggest that STAT1 plays differential roles in the coordination of CR6-specific CD4^+^ and CD8^+^ T cells, specifying CD4^+^ quality and limiting CD8^+^ quantity.

### MNV CR6-induced disease is independent of microbiome disruption in Stat1^KO^ mice

The clinical and immunological symptoms of CR6-infection in *Stat1*^KO^ mice encompass weight loss, splenomegaly, liver and spleen histopathology (Figure 1D, G-J), IL-17A production from CD4^+^ T cells (Figure 2G), as well as accumulation of neutrophils in the spleen and colon (Figure 3A), increased colonic antimicrobial gene expression (Figure S3), and increased circulating levels of pro-inflammatory cytokines, including IL-17A, IL-22, IFN*γ* and TNF*α* (Figure 3B-E). Collectively, these data suggested that microbial translocation from the intestine could be contributing to virus-induced weight loss in CR6-infected *Stat1*^KO^ mice. To address this, we isolated DNA from the liver and spleen of naïve and CR6-infected *Stat1*^Het^ and *Stat1*^KO^ mice and performed qPCR targeting the bacterial 16S gene. Notably, 16S rDNA could be detected in spleens and livers of some CR6-infected *Stat1*^KO^ mice (Figure 3F). Although neither the *α*- or *β*-diversity of fecal microbial communities was significantly affected by CR6 infection in either *Stat1*^Het^ or *Stat1*^KO^ mice (Figure 3G, H), we observed a significant infection-induced difference in the colon-associated microbial populations between *Stat1^KO^* and *Stat1^Het^* mice (Figure 3H, Table S1). These data are consistent with a possible model where virus-host gene interactions result in alterations in colon-associated microbial communities that gain access to the circulation and/or tissues and then contribute to the observed immune dysfunction and the manifestation of clinical symptoms in some *Stat1*^KO^ mice.

**Figure 3:**
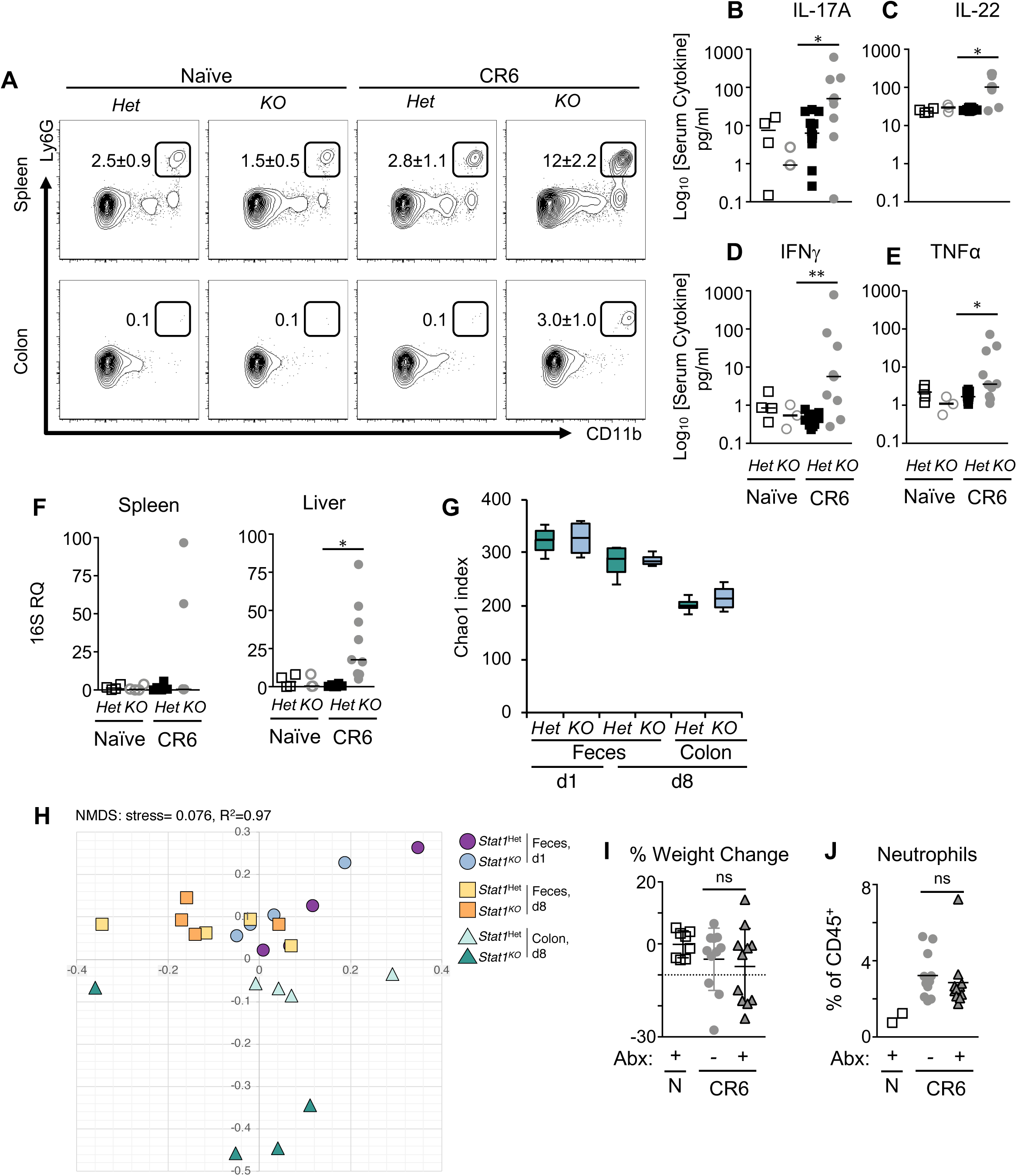
MNV CR6-induced disease is independent of microbiome disruption in *Stat1*^KO^ mice. (A-G) *Stat1^het^* and *Stat1^KO^* mice received 10^4^ pfu MNV CR6 or PBS po and were analyzed on day 8 pi. (A) Flow cytometric detection of Ly6G^+^ CD11b^+^ neutrophils in the spleen (top) and colonic LPLs (bottom). Gated on CD45^+^ B220^-^ CD11c^-^ MHCII^-^ cells, numbers in flow plots represent frequencies ± SEM. Data are representative of two independent experiments with n=4-6 mice per group. (B-E) Serum cytokine concentrations as measured by cytometric bead array. Data are compiled from three experiments n=3-4 per naïve group and n=8-10 per infected group. (F) 16S genome copies were quantified by RT-PCR in the spleen and liver. Data is expressed as fold-change relative to *Stat1*^Het^ values and normalized to expression of *Hprt*. RQ = relative quantity. Data are compiled from 4 independent experiments with n=3-4 per naïve group, and n=6 *Stat1*^Het^, n=10 *Stat1*^KO^ CR6-infected group. (G, H) Chao1 index and NMDS plot of 16S sequencing data from feces or colonic tissue at day 1 or day 8 pi. One independent experiment with repeated sampling from n=4 CR6-infected *Stat1*^Het^ and *Stat1*^KO^ mice. (I, J) *Stat1*^KO^ mice were provided water with Splenda (control) or a cocktail of antibiotics for two weeks prior to and throughout MNV CR6 infection. Weight change as a percentage difference from initial (I) and frequency of CD45^+^ B220^-^ CD11c^-^ MHCII^-^ Ly6G^+^ CD11b^+^ neutrophils in the spleen (J) in *Stat1^KO^* mice at day 8 pi. Data are compiled from 2 independent experiments n=2 naïve and n=10-11 per infected group. Two-tailed Mann-Whitney U test comparing *Stat1*^Het^ and *Stat1*^KO^ mice was performed unless otherwise noted. \**P*<0.05, \*\**P*<0.01, ***P<0.001, \*\*\*\**P*<0.0001.

To assess whether virus-induced disease in the context of STAT1-deficiency was related to bacterial translocation, we pre-treated *Stat1*^KO^ mice with a broad-spectrum cocktail of antibiotics (ABX) that has been shown to reduce the intestinal biomass^40^. However, there was no difference in the frequency of *Stat1*^KO^ mice demonstrating symptoms of CR6-induced disease between water and ABX-treated groups (Figure 3I), and mice displaying clinical symptoms in the ABX-treated group still demonstrated neutrophilia, splenomegaly, and histopathologic changes in liver and spleen (Figure 3J and data not shown). Collectively, these data suggest that despite the correlation between CR6-induced disease in *Stat1*^KO^ mice and indicators of bacterial translocation, it is likely secondary to immunopathology or direct viral-mediated injury.

### Inappropriate Th17 skewing in Stat1^KO^ mice is CD4^+^ T cell intrinsic but dispensable for disease

STAT1 plays a key role in allowing CD4^+^ T cells to receive signals for Th1 differentiation during T cell activation^41^. Thus, the observed Th17 differentiation of STAT1-deficient T cells could be due to cell intrinsic factors, and not microbial translocation from the intestine. To determine whether STAT1 plays a cell intrinsic role in CD4^+^ Th1 cell differentiation, enriched *Stat1*^Het^ and *Stat1*^KO^ CD4^+^ T cells were stimulated *in vitro* under neutral (*α*CD3/*α*CD28) or Th1-polarizing (*α*CD3/*α*CD28 + *α*IL-4, IL-2 and IL-12) conditions for 4 days. Under these conditions, there was no impairment in *Stat1*^KO^ CD4 T cell proliferation (Figure 4A). In both neutral and Th1-polarizing conditions, *Stat1*^Het^ CD4^+^ T cells produced TNF*α* and IFN*γ* (Figure 4B). In contrast, under neutral conditions *Stat1*^KO^ mice produced elevated levels of the Th17-associated cytokines IL-17A, IL-17F and IL-22 relative to *Stat1*^Het^ (Figure 4C). Although reduced, these cytokines were still detectable under Th1-polarizing conditions and the quantity of IFN*γ* was reduced in *Stat1*^KO^ cultures compared to *Stat1*^Het^ cultures (Figure 4B, C). These data are consistent with the Th17 signatures observed in CR6-infected *Stat1*^KO^ mice.

**Figure 4:**
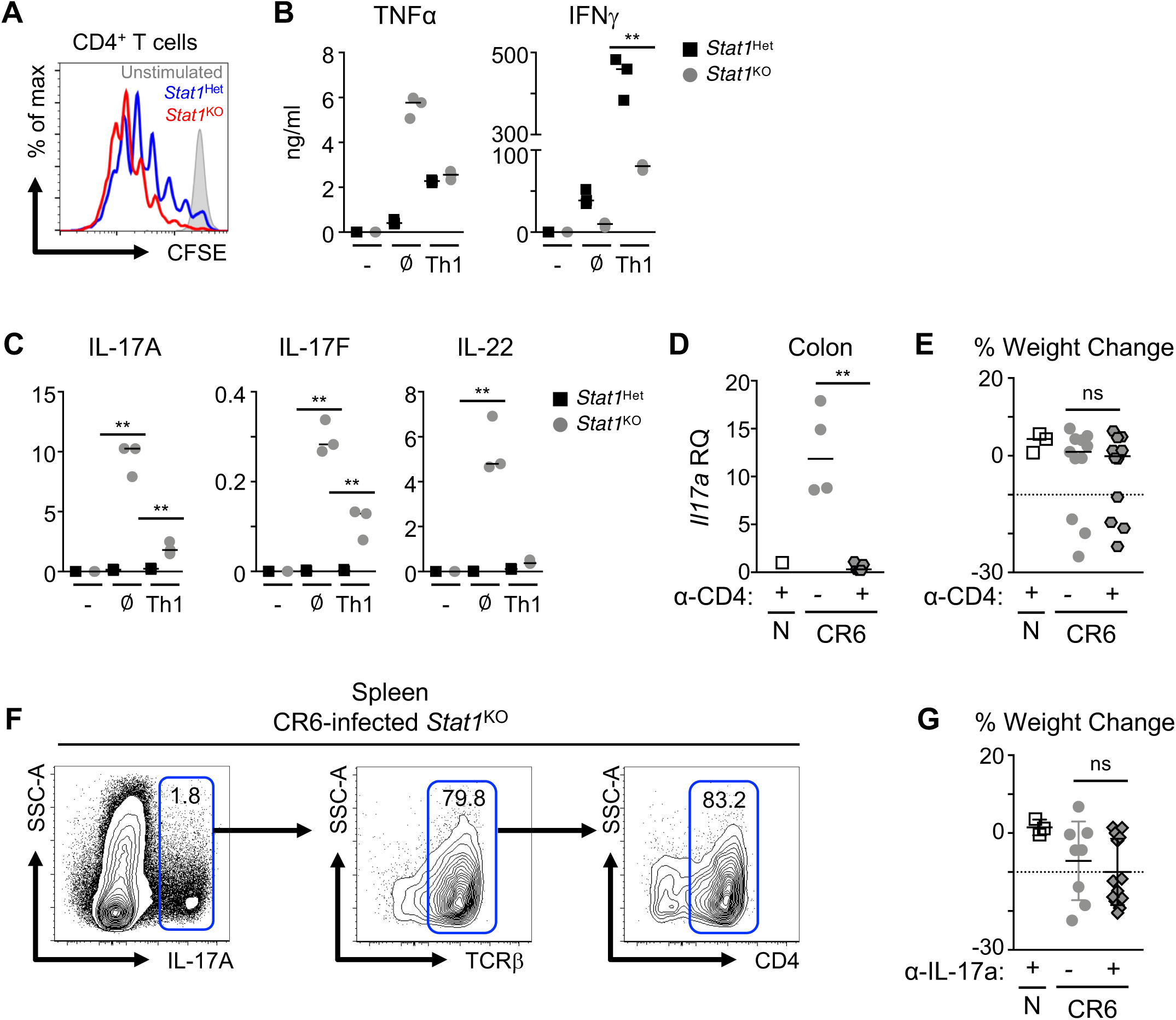
Inappropriate Th17 skewing in *Stat1*^KO^ mice is CD4^+^ T cell intrinsic but dispensable for disease. (A-C) Naïve CD4^+^ T cells were isolated from the spleens of *Stat1^het^* and *Stat1^KO^* mice and stimulated *in vitro* for 4 days with α-CD3 and α-CD28 mAbs in neutral (Ø) or Th1-skewing conditions. (A) CFSE intensity of CD4^+^ T cells stimulated in neutral conditions. (B, C) Concentrations of Th1-associated cytokines TNFα and IFNγ (B) and Th17-associated cytokines IL-17A, IL-17F, and IL-22 (B) in culture supernatant. (D, E) *Stat1^KO^* mice received 10^4^ pfu MNV CR6 or PBS po and were treated with isotype or α-CD4 mAb’s on days 0, 3 and 6 and analyzed on day 8 post-CR6 infection. (D) Expression of *IL-17A* in the colon of *Stat1^KO^* mice treated with α-CD4 mAb or isotype control, expressed as fold of *Stat^het^* values normalized to expression of *Hprt*. RQ = relative quantity. (E) Weight change in naïve and CR6-infected *Stat1*^KO^ mice treated with isotype or α-CD4 mAb. Data is representative (D) or compiled from (E) two independent experiments with n=8-10 mice per infected group. (F, G) *Stat1^KO^* mice received 10^4^ pfu MNV CR6 or PBS po and were treated with isotype or α-IL-17A mAb’s on days -1, 2 and 5 and analyzed on day 8 post-CR6 infection. (F) Flow cytometric detection of IL-17A-expressing splenocytes of MNV CR6-infected *Stat1^KO^* mice at day 8 pi. (G) Weight change in naïve and CR6-infected *Stat1*^KO^ mice treated with α-IL-17A mAb or isotype control at day 8 pi. Data is compiled from two independent experiments, n=3 naïve and n=8-12 per infected group. Two-tailed Mann-Whitney U test, \**P*<0.05, \*\**P*<0.01, \*\*\**P*<0.001, \*\*\*\**P*<0.0001.

CR6-infected *Stat1*^KO^ mice demonstrated elevated levels of serum IL-17A (Figure 3B), and increased Th17 CD4^+^ T cells compared to STAT1-sufficient controls (Figure 2G). In both viral and parasitic infections, IL-17A is pathogenic and is associated with morbidity and mortality^42, 43^. Therefore, we used antibody-mediated CD4^+^ cell depletion to test whether the dysregulated Th17 response was causing immunopathology. At day 8 post-CR6 infection, *Stat1*^KO^ mice treated with *α*-CD4 had an absence of CD4^+^ T cells in spleen and colonic compartments (Figure S4A-C), and *il17a* gene expression levels were reduced in the colon (Figure 4D), confirming efficacy of the mAb treatment. However, there was no detectable effect on CR6-induced disease due to CD4^+^ cell depletion in *Stat1*^KO^ mice (Figure 4E). Although the majority of IL-17A expressing cells are CD4^+^ T cells (Figure 4F), we postulated that other IL-17A producing cells could also contribute to disease. To test this, we treated *Stat1*^KO^ mice with either isotype or neutralizing *α*-IL-17A mAb. Again, no difference in clinical manifestation was seen between isotype or mAb-treated CR6-infected *Stat1*^KO^ animals (Figure 4G). These data are consistent with previous observations that STAT1 has a CD4^+^ T cell intrinsic role in regulating Th1 differentiation^41^ and that an inappropriate Th17 response can be generated in response to viral infection of STAT1-deficient mice^43^. In the context of a LCMV Armstrong, a viral pathogen that induces a robust type I IFN response, dysregulated, pro-inflammatory CD4^+^ T cells mediate virus-induced mortality^25^. However, in the context of this normally asymptomatic, commensal-like intestinal virus, the dysregulated CD4^+^ T cell response caused by STAT1-deficiency does not have a direct role in viral-induced pathogenesis.

### CD8^+^ T cells do not mediate MNV CR6-induced disease in Stat1^KO^ mice

Previous studies have shown that failure to regulate the magnitude of a CD8^+^ T cell response can contribute to immunopathology^44–46^. *Stat1*^KO^ mice infected with CR6 have significantly expanded populations of MNV-specific CD8^+^ T cells, increased numbers of IFN*γ* and TNF*α*-producing CD8^+^ T cells (Figure 2A-C) and elevated levels of circulating IFN*γ* and TNF*α* (Figure 3D, E). To determine whether this dysregulated population of virus-specific CD8^+^ T cells could contribute to immunopathology, we depleted CD8α^+^ cells and assessed clinical scores and viral loads in *Stat1*^KO^ mice. Despite efficient depletion of CD8^+^ cells (Figure S4D-F), there was no significant difference in virus-induced weight loss between isotype and *α*-CD8*α* mAb treated *Stat1*^KO^ mice (Figure 5A). However, there was a loss of neutrophil accumulation in the spleens of *α*-CD8 mAb treated *Stat1*^KO^ mice (Figure 5B), providing indirect evidence that CR6-induced disease is unrelated to neutrophilia. Notably, depletion of CD8^+^ T cells had no impact on either intestinal or splenic viral burdens (Figure 5C), indicating that loss of antiviral effector cells does not exacerbate impaired control of viral replication.

**Figure 5:**
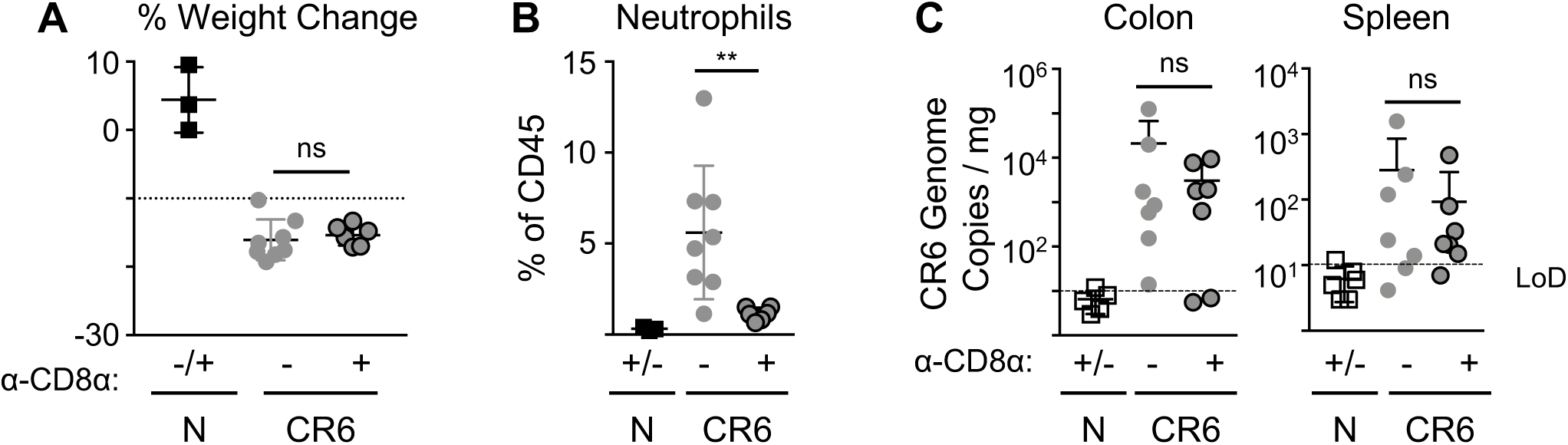
CD8^+^ T cells do not mediate MNV CR6-induced disease in *Stat1*^KO^ mice. *Stat1^KO^* mice received 10^4^ pfu MNV CR6 or PBS iv and were treated with isotype or α-CD8 mAb’s on days 0, 3 and 6 and analyzed on day 8 post-CR6 infection. (A) Weight change in naïve and CR6-infected *Stat1*^KO^ mice treated with α-CD8 mAb or isotype control at day 8 pi. (B) Frequency of CD45.2^+^ B220^-^ CD11c^-^ MHCII^-^ Ly6G^+^ CD11b^+^ neutrophils in the spleen. (C) CR6 genome copies in colon (left) and spleen (right) were quantified by RT-PCR. Data are from one independent experiment, n=3 naïve and n=7-8 per infected group. Mann-Whitney U test. \**P*<0.05, \*\**P*<0.01, ***P<0.001, \*\*\*\**P*<0.0001. LoD: Limit of Detection.

### Clinical symptoms of viral-induced disease and immune dysregulation can be uncoupled by antiviral treatment of Stat1^KO^ mice

In the absence of T cell mediated immunopathology or microbial translocation causing disease, we investigated whether direct inhibition of viral replication was sufficient to prevent disease in CR6-infected *Stat1*^KO^ mice. To this end, we treated mice with 2’-C-methylcytidine (2CMC), a viral polymerase inhibitor previously shown to limit MNV replication^47^, starting concurrently with CR6 infection (Figure 6A). Consistent with previous reports, this prophylactic treatment regimen resulted in viral burdens at or below the limit of detection in both colon and spleen (Figure 6B) and prevented virus-induced weight loss (Figure 6C). In order to allow early stages of infection to proceed, we used a modified treatment regimen and started treatment with 2CMC at day 3 post-infection (Figure 6D). Therapeutic treatment with 2CMC was associated with significantly reduced viral burdens. In the colon, burdens of 2CMC-treated mice were at or near the limit of detection and there was a reduction in detectable genome copies in the spleens of 2CMC-treated animals compared to PBS-treated control *Stat1*^KO^ mice (Figure 6E). Notably, at day 8 pi, therapeutic 2CMC treatment prevented CR6-induced weight loss in *Stat1*^KO^ mice (Figure 6F). Consistent with the lack of clinical signs, therapeutic 2CMC treatment prevented CR6-induced splenomegaly, liver and spleen necrosis associated with CR6-infection of *Stat1*^KO^ mice (data not shown). Finally, we addressed whether 2CMC treatment also impacted the dysregulated antiviral CD4^+^ or CD8^+^ T cell responses that are associated with CR6-induced disease in *Stat1*^KO^ mice. Notably, despite significant reduction in viral burdens, the frequency and cell numbers of CD8^+^ MNV-specific T cells, ROR*γ*t^+^ CD4^+^ Th17 cells, and neutrophils were equivalent between vehicle and 2CMC-treated *Stat1*^KO^ mice (Figure 6G-J). Thus, our data uncouple impaired viral control from STAT1-directed immune cell polarization and function, suggesting that neither immunopathology nor secondary bacterial translocation are directly involved in viral-induced disease. Instead, STAT1-dependent tolerance is maintained by restricting the virus to mucosal tissues and limiting damage to systemic organs caused by unchecked viral replication.

**Figure 6:**
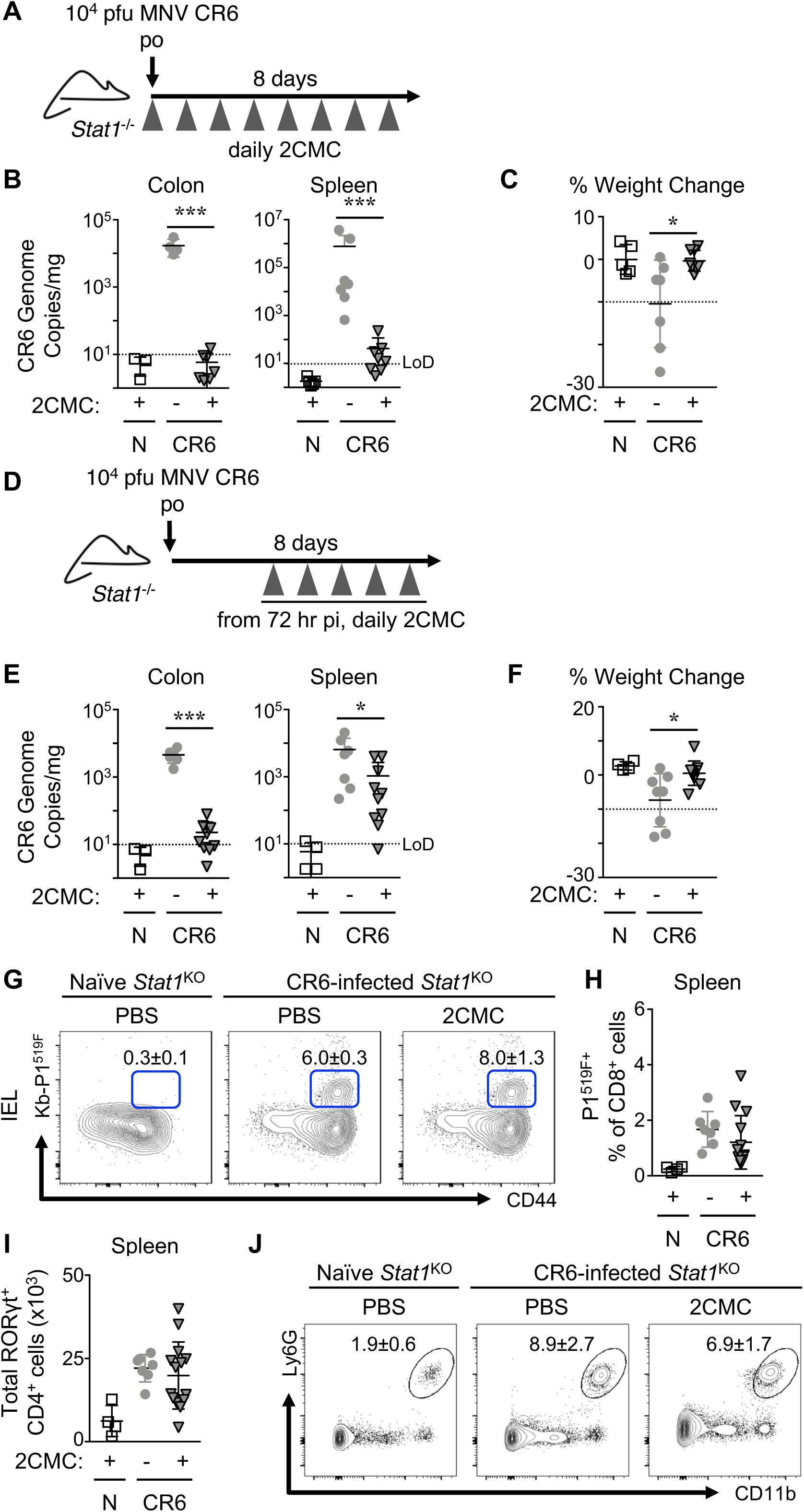
Clinical symptoms of viral-induced disease and immune dysregulation can be uncoupled by antiviral treatment of *Stat1*^KO^ mice. (A) Experimental scheme for early treatment regimen used in (B, C). Data are compiled from two independent experiments, n=6 naïve and n=7-8 per infected group. (B) CR6 genome copies in colon and spleen were quantified by RT-PCR. (C) Weight change as a percentage difference from initial. (D) Experimental scheme for late treatment regimen used in (E-J). Data are compiled from two independent experiments, n=4 naïve and n=7-8 per infected group. (E) CR6 genome copies in colon and spleen were quantified by RT-PCR. (F) Weight change as a percentage difference from initial. (G) Flow cytometric detection of splenic MNV-CR6 P1^519F^ antigen-specific CD8^+^ T cells isolated from colonic IELs. Numbers in flow plots represent frequencies ± SEM of CR6 Kb-P1^519F^ tetramer^+^ cells within the CD8^+^ lymphocyte gate. (B) Quantification of MNV CR6-specific cells as a frequency of CD45^+^ CD8^+^ splenocytes. (I) Total RORγT^+^ CD4^+^ T cells in the spleen. (J) Flow cytometric detection of Ly6G^+^ CD11b^+^ neutrophils in colonic LPLs. Gated on CD45^+^ B220^-^ CD11c^-^ MHCII^-^ cells, numbers in flow plots represent frequencies ± SEM. Mann-Whitney U Test. \**P*<0.05, \*\**P*<0.01, ***P<0.001, \*\*\*\**P*<0.0001. LoD: Limit of Detection.

## Discussion

Recent work demonstrating the ability of the virome to both positively and negatively influence our health has underscored the importance of elucidating how these relationships are regulated by the host. The mammalian virome includes viruses that have evolved distinct strategies for long-term persistence, such as latency, genomic integration and/or immune evasion. In both mice and humans, Noroviruses can establish long-term persistence without latency. Experimental infection of volunteers has demonstrated that human norovirus (huNoV) can establish persistent infection for weeks or months after resolution of clinical symptoms, and despite high viral loads some patients never experience clinical symptoms^48^. These results, together with the demonstration that MNV can confer host-protective effects in mice^7, 10^, suggest that norovirus infection fits the profile of mutualism; however, the host mechanisms facilitating tolerance of this infection are not well understood^10, 49^. Here, we describe profound immune dysregulation following MNV CR6 infection of STAT1-deficient mice. However, unlike other pathogenic viral infections where immune-mediated pathology contributed to morbidity and mortality, we demonstrate that controlling viral replication via treatment with a viral polymerase inhibitor is sufficient to abrogate disease even in the presence of immune dysfunction. These data uncouple the STAT1-mediated effects on antiviral T cell differentiation and function from those required for direct inhibition of viral replication.

In WT mice, CR6 infection is limited to the colonic epithelium, where it infects tuft cells expressing the viral entry receptor, CD300lf^18^. However, we detected high viral burdens in multiple peripheral tissues, including the spleen and liver. Previous studies suggest that phagocytes, including monocytes and neutrophils, express CD300lf and that some myeloid subsets can be recruited to the intestine in an IFNAR-independent manner^10, 50^. Further, loss of STAT1 signaling in CD11c^+^ dendritic cells permits systemic and persistent replication of MNV CW3^51^. Thus, we hypothesize that the systemic dissemination of CR6 in *Stat1*^KO^ mice could be mediated by migratory phagocytes that allow for continued spread into tissue resident cells that are permissive for CR6 infection. The failure to control CR6 replication in tissues could then lead to organ failure and underlie the observed mortality in *Stat1*^KO^ mice.

Consistent with a previous report^52^, CR6 infection had no effect on luminal or colon-associated microbial communities in STAT1-sufficient animals. However, maintenance of colonic-tissue associated community structure was STAT1-dependent, which we considered indicative of potential bacterial translocation that could contribute to CR6-induced pathology. However, systemic bacterial dissemination did not correlate with clinical signs and depletion of bacterial biomass had no effect on CR6-induced disease in *Stat1*^KO^ mice. While these results do not eliminate the possibility that the microbiota contribute to the observed disease, they suggest that dysbiosis is secondary to virus-induced morbidity.

Previous reports have also documented infrequent mortality of B6/*Stat1*^KO^ mice upon infection with CR6^53^. Strains MNV-O7 and MNV4 form persistent infections similar to MNV CR6 in wild-type mice, but neither cause mortality in *Stat1* ^KO^ mice^33, 34^. Furthermore, MNV4 induces colonic inflammation in *Stat1*^KO^ mice, while in our facility colonic architecture was maintained in CR6-infected *Stat1*^KO^ mice. These data indicate that strain-specific differences may provide useful insights into viral determinants of tolerance. Notably, CR6 infection of *Stat1*^KO^ mice significantly up-regulated expression of epithelial-protective factors such as IL-22 and the antimicrobial peptides Reg3*β* and Reg3*γ*, which may contribute to maintenance of intestinal homeostasis. In the context of pathogenic RV infection, IL-22 and IFN*λ* are reported to synergize in a STAT1-dependent manner to up-regulate interferon stimulated genes (ISGs) that limit viral replication in enterocytes and thus protect the intestinal epithelial barrier^22^. Although viral loads are higher in the colon of *Stat1*^KO^ mice compared to littermate controls, CR6 epithelial tropism is limited to rare tuft cells and the IL-22/STAT1 convergent activation of ISGs may be less important in this context. However, CR6 colonization fortifies the intestinal barrier of WT GF mice via IL-22-mediated STAT3 activation and up-regulation of Reg3*β*^10^ and this pathway may limit intestinal damage in CR6 colonized *Stat1*^KO^ mice.

STAT1 transduces signals from type I, II, and III interferons, and previous reports have investigated the individual roles of these signals in the context of CR6. Despite only weakly inducing type I interferon, CR6 infection of *Ifnar^-/-^* mice leads to systemic viral dissemination. In contrast, loss of type III interferon signaling leads to increased viral shedding, but not dissemination, while loss of type II interferon signaling has no documented effect on CR6 infection^20^. Finally, combined deficiency of type I and II IFN signaling results in increased viral shedding^47^. Notably, the loss of any one of these signaling pathways in isolation does not lead to increased morbidity and mortality; however, we hypothesize that in the case of *Stat1*^KO^ mice, the combined loss of type I and type III signaling leads to systemic dissemination of a higher load of virus from the colon, resulting in a loss of control of systemic viral replication and eventual morbidity and mortality.

Our data demonstrate the novel finding that 2CMC remains efficacious against CR6 when administered therapeutically in a severely immunocompromised model; given the risks posed by Noroviruses to immunocompromised humans, this is further evidence for the potential usefulness of targeted antiviral treatment in the clinic. Additionally, STAT1 suppression is an off-target effect of the purine analog fludarabine, a chemotherapeutic agent used in the treatment of leukemia and lymphoma^54^. Patients treated with fludarabine are at an elevated risk of opportunistic viral infection, and our data supports the hypothesis that that fludarabine-mediated suppression of STAT1 may contribute to this risk of infection. Based on this, treatment with fludarabine may result in a loss of tolerance to ongoing - and often sub-clinical - norovirus infections, potentially increasing the risk of complications during treatment. Together, our data highlight the importance of STAT1 for maintaining tolerance to chronic viral infections.

## Materials and Methods

### Mouse strains and housing conditions

Mice were housed at the University of British Columbia (UBC) specific pathogen-free Centre for Disease Modeling. MNV-free C57BL/6 *Stat1^+/-^* mice (The Jackson Laboratory, strain B6.129S(Cg)-*Stat1^tm1Dlv^/J* (012606) were bred as either *Stat1^+/-^* x *Stat1^+/-^* or *Stat1^+/-^* x *Stat1^-/-^*. To control for microbiome and other environmental effects^55^, resulting *Stat1*^WT^, *Stat1*^Het^ and *Stat1*^KO^ littermates were used. If an individual experiment required multiple litters, age-matched STAT1- sufficient and -deficient mice from each litter were included. Mixed genotypes were co-housed before and throughout the experiment. Replicate experiments were performed on litters from multiple dams. Experimental mice were aged-matched within each experiment, and were 6-12 weeks of age for all experiments.

All experiments were performed according to guidelines from UBC Animal Care Committee and Biosafety Committee-approved protocols. Mice were housed in ventilated Ehret cages prepared with BetaChip bedding and had *ad libitum* access to irradiated PicoLab Diet 5053 and reverse osmosis/chlorinated (2-3 ppm)-purified water. Housing rooms were maintained on a 14/10-hour light/dark cycle with temperature and humidity ranges of 20-22°C and 40-70%, respectively. Sentinel mice housed in experimental rooms were maintained on dirty bedding and nesting material and were tested on a quarterly basis for presence of mites (*Myocoptes, Radford/Myobia)*, pinworm (*Aspiculuris tetaptera, Syphacia obvelata*), fungi (*Encephalitozooan cuniculi*), bacteria (*Helicobacter* spp., *Clostridium piliforme, Mycoplasma pulmonis,* CAR Bacillus), and viruses (Ectromelia, EDIM/Rotavirus, MHV, MNV, MPV, MVM, LCMV, MAV1/2, MCMV, Polyoma, PVM, REO3, Sendai and TMEV).

### Infection, monoclonal antibody, antibiotic and antiviral treatments

Mice were infected by oral gavage (per os, po) or intravenously (iv) by tail vein injection with 10^4^ pfu of MNV CR6 diluted in sterile PBS. Naive mice were sham-infected with sterile PBS (Sigma). Mice were monitored every 24-48 hours and euthanized at humane endpoint (loss of ≥20% of initial body weight).

To deplete CD4^+^ or CD8^+^ cells, 400 μg of either *α*-CD4 (clone GK1.5, UBC AbLab) or *α*-CD8*α* antibodies (clone 53.67, UBC AbLab) was administered intraperitoneally (ip) at days 0, 3 and 6 post-MNV CR6 infection. Cellular depletion in spleens and intraepithelial lymphocyte compartments was monitored by flow cytometric detection with *α*-CD4 BV650 (clone RM4-5, BD Biosciences) or *α*-CD8*β* (clone H35-17.2, ThermoScientific). To neutralize IL-17A, 500 μg *α*-IL-17A (clone 17F3, BioXcell) was administered ip at days -1, 2 and 5 of MNV CR6 infection. For all mAb treatment experiments, antibodies were prepared in sterile PBS (Sigma) and IgG1 isotype control antibodies (clone MOPC-21, BioXCell) were administered with the same dosing regimen and schedule as the treatment.

For antibiotic administration, water supplemented with 0.25 mg/ml vancomycin, 0.5 mg/ml neomycin, 0.5 mg/ml gentamicin, 0.5 mg/ml ampicillin, 0.25 mg/ml metronidazole and 4 mg/ml Splenda was provided *ad libitum*^40^. Control mice received sterile drinking water supplemented with 4 mg/ml Splenda.

### Tissue histology

Samples were flushed with PBS and fixed in 4% paraformaldehyde and embedded in paraffin. 5 µm sections were cut and used for hematoxylin & eosin staining (Histochemistry Service Laboratory at the University of British Columbia). Histology images were acquired with Zeiss Axio Observer 7 and an AxioCam 105 microscope camera.

### Statistical analysis

Statistical analyses were performed using GraphPad Prism (GraphPad Software, Version 6.0). Two-tailed nonparametric Mann-Whitney U-test, Mantel-Cox or Spearman correlation coefficient, as appropriate (specified in figure captions). Values are reported as mean ± SEM. * p < 0.05, ** p < 0.01, *** p < 0.001, **** p < 0.0001.

## Acknowledgements

We thank members of the Osborne laboratory for discussions and critical reading of the manuscript. Research in the Osborne lab is supported by the Natural Sciences and Engineering Research Council of Canada, the Canadian Institutes of Health Research, the Canada Research Chair program and scholarships from the University of British Columbia. The authors declare no potential conflicts of interest. The authors would like to thank the NIH Tetramer Core Facility for providing MHCI (Kb-P1^519Y^ and Kb-P1^519F^) and MHCII (IAb-P1^496^) tetramers. The authors would like to thank the University of British Columbia Flow Cytometry Core and the Centre for Disease Modeling for their assistance and support.

## Author Contributions

HAF & LCO conceived the study. HAF, AJS, NMF, WY, BKH, HGR & JHS performed *in vivo* and *in vitro* experiments and analyzed data. RLS & SAC performed bioinformatic analysis of bacterial community composition. JR-P & JN contributed 2CMC and valuable insight for experimental design and interpretation. IW analyzed tissue histology. AJS & LCO prepared the manuscript.

## Supplemental information

### Supplemental Figure Legends

**Figure S1:**
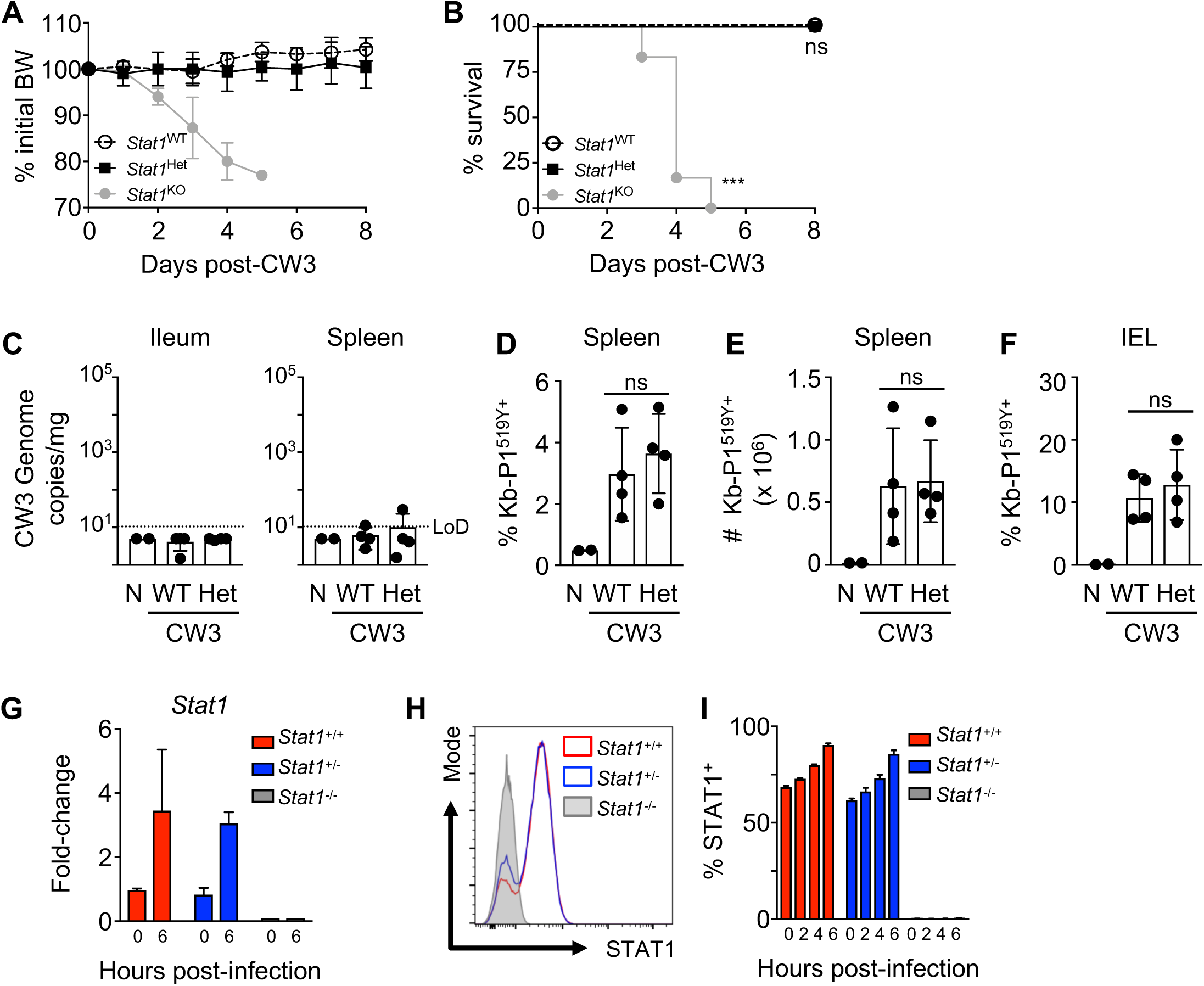
*Stat^Het^* phenocopy *Stat1^WT^* mice. (A-F) *Stat1^Het^* and *Stat1^WT^* mice were infected with 10^4^ pfu MNV CW3 or PBS po and euthanized on day 8 or at humane endpoint. Data are representative of two independent experiments with n=3-4 mice per group. (A) Weight change as a percentage difference from initial. (B) Survival curve of CW3-infected mice, Mantel-Cox Test. (C) CW3 genome copies in ileal and splenic tissue in naïve and infected *Stat1^WT^* and *Stat1^Het^* mice were quantified by RT-PCR. (D-F) Frequency (D) and total number (E) of MNV-CW3 P1^519Y^ antigen-specific CD8^+^ T cells isolated from the spleen and small intestinal IEL (F). Data are from one independent experiment. (G-I) BMDCs derived from *Stat1*^WT^*, Stat1*^Het^, and *Stat1*^KO^ mice were pulsed for one hour with MNV CW3 (MOI = 0.5), washed and analyzed at the indicated time. Data are compiled from 3 independent experiments with triplicate samples from individual donors. LoD: Limit of Detection. (G) *Stat1* gene expression, normalized to *Hprt* in *Stat1*^WT^ cells. (H) Flow cytometric detection of STAT1 in BMDCs at 6 hours pi. (I) Kinetics of MNV CW3 infection-induced STAT1 expression in BMDCs.

**Figure S2:**
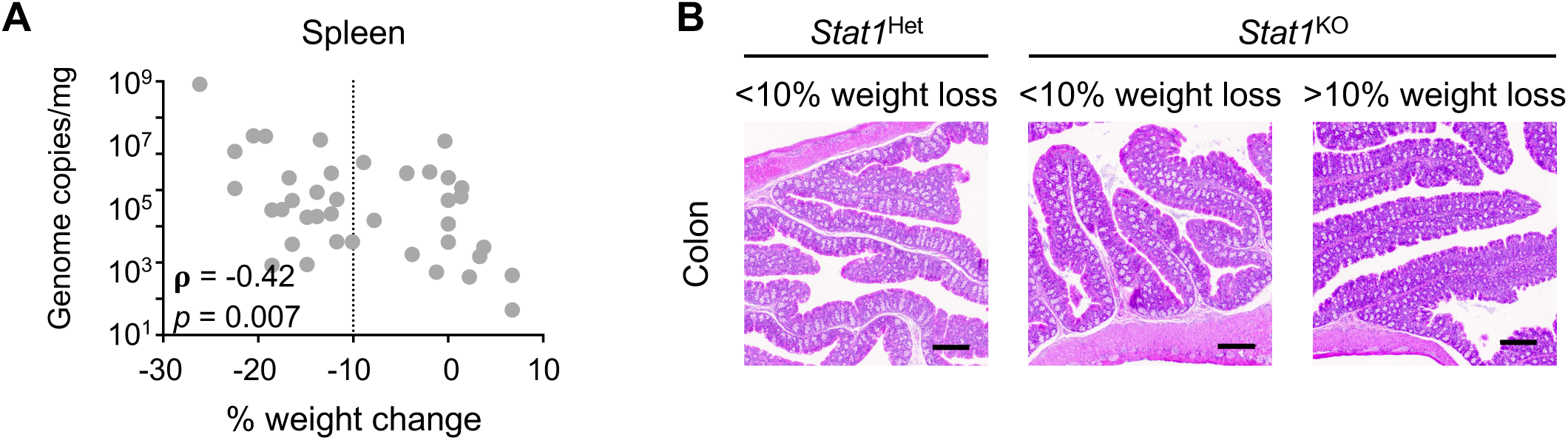
STAT1-dependent tolerance of MNV CR6 infection. *Stat1^het^* and *Stat1^KO^* mice received 10^4^ pfu MNV CR6 or PBS po and were analyzed on day 8 pi. (A) Correlation between MNV CR6 genome copies in spleen and weight loss in *Stat1^KO^* mice, Spearman Correlation Coefficient Test. (B) Representative H&E stained colon sections at day 8 pi, scale bar = 200 µM. Data is representative of three independent experiments.

**Figure S3:**
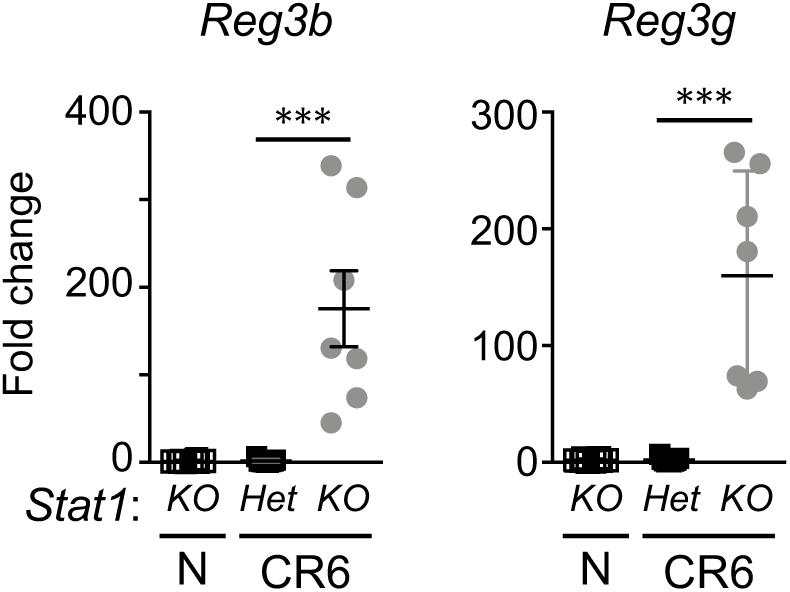
STAT1 controls expression of intestinal antimicrobial peptides. *Stat1^het^* and *Stat1^KO^* mice received 10^4^ pfu MNV CR6 or PBS po and were analyzed on day 8 pi. Expression of *Reg3b* and *Reg3g* expressed as fold of naïve *Stat1^WT^* values normalized to expression of *Hprt*. Mann-Whitney U Test. Data are representative of 3 independent experiments, n=7 per group.

**Figure S4:**
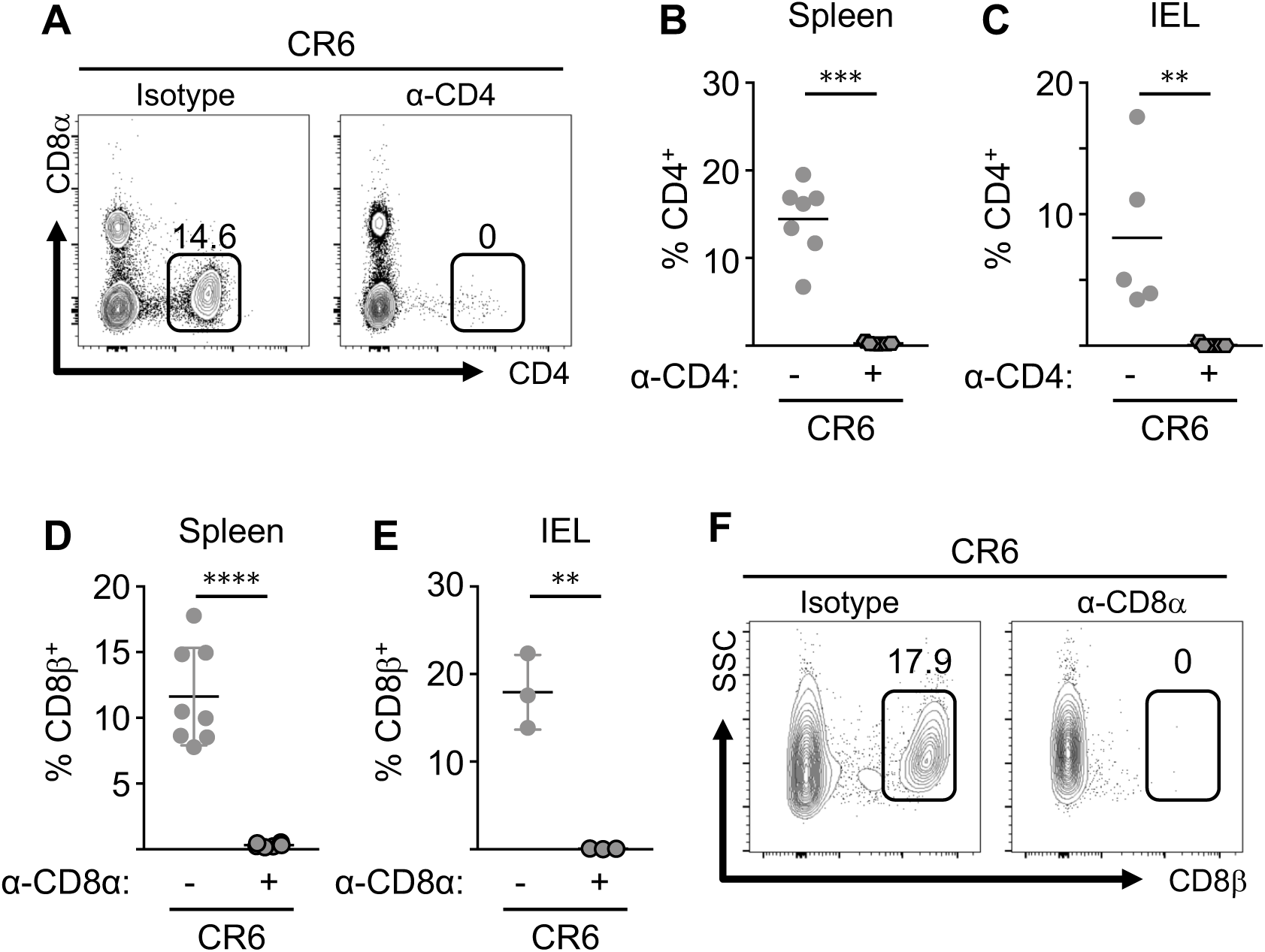
Antibody-mediated depletion of CD4^+^ and CD8^+^ cells in the spleen and colon. *Stat1^KO^* mice received 10^4^ pfu MNV CR6 or PBS po and were treated with α-CD4 (A-C) or α-CD8 (D-F) mAb or isotype control ip on days 0, 3, and 6 pi and analyzed on day 8 pi. (A) Representative staining of CD4^+^ splenocytes. (B) Frequency of CD4^+^ cells as a percentage of CD45^+^ splenocytes. (C) Frequency of CD4^+^ IELs as a percentage of TCRβ^+^ splenocytes. Data from (A-C) are compiled from two independent experiments with a minimum of n=5 per group. (D, E) Frequency of CD8^+^ cells as a percentage of CD45^+^ splenocytes (d) and IELs (E). (F) Representative staining of CD8^+^ IELs. Data from (D-F) are from one independent experiment with n=7-8 mice per group.

### Supplementary Materials and Methods

#### Flow cytometry

Directly conjugated antibodies used for surface and intracellular cytokine staining were obtained from BioLegend: CD8*α* PE-Cy7 (53-6.7), CD44 APC-Cy7 (IM7), TCR*β* PerCP-Cy5.5 and PE/Dazzle 594 (H57-597), CD45.2 AlexaFluor 700 (104), TNF*α* PE (MP6-XT22), IL-17A PE/Dazzle 594 (TC11-18H10.1), T-bet Brilliant Violet 421 (4B10); BD Biosciences: CD4 BV650 (RM4-5); or eBioscience/ThermoFisher: MHCII I-A/I-E APC (M5/114.15.2), IFN*γ* eFluor450 (XMG1.2), Foxp3 Alexa Fluor 488 (FJK-16s), ROR*γ*t PerCP-eFluor 71 (B2D). Tetramers specific for MNV CR6 capsid (MHCI Kb-P1^519F^ and MHCII IA^b^-P1^496^) were generated by the NIH Tetramer Core Facility ^38^. Samples were collected on an LSR II (BD Bioscience) using FACS Diva software. Data files were analyzed using FlowJo software (version 10.1r3, Tree Star Inc.). For all analyses, dead cells that stained positive for LIVE/DEAD Fixable Aqua Dead Cell Stain (Invitrogen, Molecular Probes) were excluded.

#### Supernatant and serum cytokine detection

For *in vitro* CD4^+^ T cell activation, cells were isolated from combined spleens and mesenteric lymph nodes of *Stat1*^Het^ or *Stat1*^KO^ mice. CD4^+^ T cells were enriched for using the StemCell Untouched CD4 T cell isolation kit (STEMCELL Technologies). Purity was typically >90% CD45^+^ TCRβ^+^ CD4^+^ cells as frequency of total live cells, as measured by flow cytometry. Following enrichment, 10^5^ cells were plated in RPMI-1610 supplemented with 10% fetal bovine serum, 2µM L-glutamine, 50U/mL penicillin/streptomycin, 25mM HEPES, and 55µM β-mercaptoethanol. Cells were then given no stimulation, neutral stimulation (1 μg/ml each platebound *α*-CD3, soluble *α*-CD28,) or Th1-skewing stimulation (*α*-CD3, *α*-CD28, 10 μg/ml *α*-IL-4, 10 ng/ml IL-2, 10 ng/ml IL-12) for 4 days. Supernatant was harvested and stored at −20°C for further processing.

Concentrations of serum and supernatant cytokines were assessed using the Cytometric Bead Array Mouse Inflammation kit and Mouse Th Cytokine panel (BioLegend LEGENDPlex) according to manufacturer instructions. Blood was collected from euthanized mice and cleared serum recovered and stored at −80°C. Samples were run on the LSR II (BD Bioscience) using FACS Diva software and analyzed using FlowJo software. Standard curves were constructed using non-linear regression in Excel, utilizing the SOLVER plugin^64^.

#### MNV CR6 generation and quantification

Stocks of MNV CR6 were prepared as previously described^38^. Briefly, 293T cells (ATCC, Manassas, VA) were transfected with a plasmid containing the genome of MNV-1.CR6 (GenBank accession number EU004676) and FuGENE-HD (Promega). After 48 hrs at 37°C, transfected 293T cells were lysed by freeze-thaw cycle and centrifuged to remove cellular debris. Cleared supernatant was transferred to RAW 264.7 cells and incubated for 48 hours. After a freeze-thaw, the supernatant was cleared of debris, assayed for plaque forming units by plaque assay and stored at −80°C.

#### Quantitative PCR for gene expression and measurement of viral loads

Cells, colon, liver, or spleen were harvested into RNAlater (Ambion). RNA was isolated using PureLink RNA Mini Kit (ThermoFisher Scientific). cDNA conversion was achieved using random hexamer priming and Invitrogen Superscript III Reverse Transcriptase (ThermoFisher Scientific). Detection of MNV genome copies in tissue samples was performed as previously described on a Quant Studio 3 Real-Time PCR System (Applied Biosystems)^56^. Absolute numbers of MNV genome copies were extrapolated from a standard curve and normalized to tissue weight. Gene expression analysis was performed using PowerUp SYBR Green Master Mix (Applied Biosystems) on a Quant Studio 3 Real-Time PCR System (Applied Biosystems). Gene expression was normalized to *hprt1* and is expressed as fold-induction relative to the average expression levels in naïve *Stat1*^Het^ mice.

#### Lymphocyte recovery and ex vivo lymphocyte stimulation

Single cell suspensions of splenic lymphocytes were prepared by mashing tissue and passing cells through a 70 μm nylon mesh filter followed by ACK lysis to remove red blood cells. Intestinal lymphocytes were harvested from the cecum and colon as previously described^38^. For peptide-specific cytokine responses, 2×10^6^ lymphocytes were stimulated with 0.4 µg/ml MHC class I-restricted peptides (P1^519Y^)^38^ or 0.8 µg/ml MHC class II-restricted peptides (Nterm^28^, Nterm^46^, NTPase^622^, Pro^991^, Pol1^630^, P1^496^)^56^ for 5 hr at 37°C in the presence of 10 µg/ml brefeldin A (BFA) (Sigma) and GolgiStop (BD Biosciences). For polyclonal cytokine responses, lymphocytes were stimulated with 0.1 µg/ml PMA and 1 µg/ml ionomycin (Sigma) in the presence of BFA and GolgiStop for 5 hr at 37°C. Intracellular cytokine and effector molecule staining was performed using Cytofix/Cytoperm and Permeabilization Buffer (BD Biosciences). To detect MNV-specific CD4^+^ T cells, cells from spleen, peripheral, and mesenteric lymph nodes were pooled, passed through a 70 μm nylon mesh filter to create a single cell suspension, labeled with tetramers specific for MNV CW3 capsid P1^496^ and subjected to magnetic bead enrichment, as described in^39^. Transcription factor staining was performed using Foxp3 Transcription Factor Fixation and Permeabilization Buffer (eBioscience/ThermoFisher), according to manufacturer instructions.

For *in vitro* CD4^+^ T cell activation, cells were isolated from combined spleens and mesenteric lymph nodes of *Stat1*^Het^ or *Stat1*^KO^ mice. CD4^+^ T cells were enriched for using the StemCell Untouched CD4 T cell isolation kit (STEMCELL Technologies). Purity was typically >90% CD45^+^ TCRβ^+^ CD4^+^ cells as frequency of total live cells, as measured by flow cytometry. Following enrichment, 10^5^ cells were plated in RPMI-1610 supplemented with 10% fetal bovine serum, 2µM L-glutamine, 50U/mL penicillin/streptomycin, 25mM HEPES, and 55µM β-mercaptoethanol. Cells were then given no stimulation, neutral stimulation (1 μg/ml each platebound *α*-CD3, soluble *α*-CD28,) or Th1-skewing stimulation (*α*-CD3, *α*-CD28, 10 μg/ml *α*-IL-4, 10 ng/ml IL-2, 10 ng/ml IL-12) for 4 days. Supernatant was harvested and stored at −20°C for further processing.

#### Bacterial DNA extraction, amplification and iTag sequencing

DNA was extracted from liver, spleen, colon tissue and fecal pellets using DNeasy PowerSoil Kit (Qiagen) as per manufacturer’s instructions. Resulting DNA was stored at −20°C. Liver and spleen 16S rDNA amplification was measured via SYBR incorporation detected on a Quant Studio 3 Real-Time PCR System (Applied Biosystems). Quantification was conducted using the PicoGreen Assay (Invitrogen) for dsDNA^57^ measured on a TECAN M200 plate reader. Bacterial and archaeal SSU rRNA gene fragments from the extracted genomic DNA were amplified using primers 515F and 806R. Sample preparation for amplicon sequencing was performed as described as in ^58, 59^. The amplicon library was analyzed on an Agilent Bioanalyzer using the High Sensitivity DS DNA assay to determine approximate library fragment size, and to verify library integrity. Pooled library concentration was determined using the KAPA Library Quantification Kit for Illumina. Library pools were diluted to 4 nM and denatured into single strands using fresh 0.2 N NaOH as recommended by Illumina. The final library was loaded at a concentration of 8 pM, with an additional PhiX spike-in of 5-20%. Sequencing was conducted through the UBC Sequencing and Bioinformatics Consortium (https://sequencing.ubc.ca/).

#### Bioinformatic analysis

Sequences were processed using Mothur^60^. In brief, sequences were removed from the analysis if they contained ambiguous characters, had homopolymers longer than 8 bp and did not align to a reference alignment of the correct sequencing region. Unique sequences and their frequency in each sample were identified and then a pre-clustering algorithm was used to further de-noise sequences within each sample^61^. Unique sequences were identified and aligned against a SILVA alignment (available online at http://www.mothur.org/wiki/Silva_reference_alignment). Sequences were chimera checked using UCHIME^62^ and reads were then clustered into 97% OTUs using OptiClust^63^. OTUs were classified using the SILVA reference taxonomy database (release 132, available at http://www.mothur.org/wiki/Silva_reference_files). All data was visualized in R and Excel. For alpha and beta diversity measures, all samples were subsampled to the lowest coverage depth and standard indices were calculated in Mothur. Community structure was investigated using the Yue and Clayton similarity estimator, non-metric multidimensional scaling (NMDS)) was used to compare bacteria community structures across all samples, and a Multiple sample Analysis of Molecular Variance (AMOVA) was used to test the significance of differences between microbial communities.

---

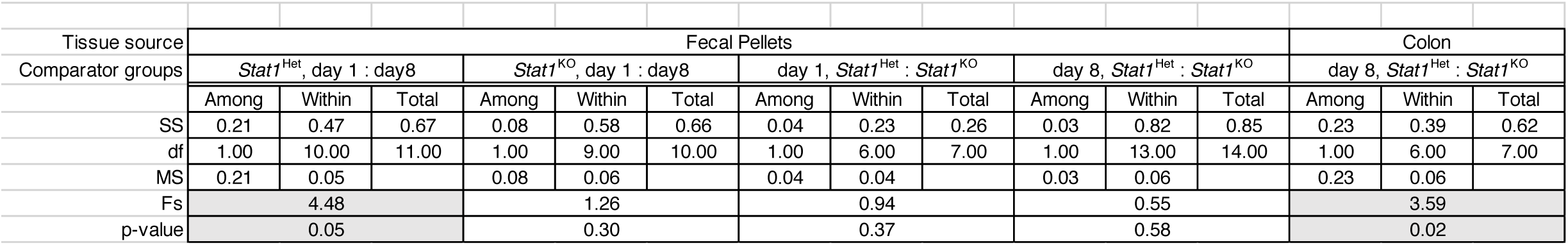

|  |  |  |  |  |  |  |  |  |  |  |  |  |  |  |  |
| --- | --- | --- | --- | --- | --- | --- | --- | --- | --- | --- | --- | --- | --- | --- | --- |
| Tissue source | Fecal Pellets |  |  |  |  |  |  |  |  |  |  |  | Colon |  |  |
| Comparator groups | *Stat1*<sup>Het</sup>, day 1 : day8 |  |  | *Stat1*<sup>KO</sup>, day 1 : day8 |  |  | day 1, *Stat1*<sup>Het</sup> : *Stat1*<sup>KO</sup> |  |  | day 8, *Stat1*<sup>Het</sup> : *Stat1*<sup>KO</sup> |  |  | day 8, *Stat1*<sup>Het</sup> : *Stat1*<sup>KO</sup> |  |  |
|  | Among | Within | Total | Among | Within | Total | Among | Within | Total | Among | Within | Total | Among | Within | Total |
| SS | 0.21 | 0.47 | 0.67 | 0.08 | 0.58 | 0.66 | 0.04 | 0.23 | 0.26 | 0.03 | 0.82 | 0.85 | 0.23 | 0.39 | 0.62 |
| df | 1.00 | 10.00 | 11.00 | 1.00 | 9.00 | 10.00 | 1.00 | 6.00 | 7.00 | 1.00 | 13.00 | 14.00 | 1.00 | 6.00 | 7.00 |
| MS | 0.21 | 0.05 |  | 0.08 | 0.06 |  | 0.04 | 0.04 |  | 0.03 | 0.06 |  | 0.23 | 0.06 |  |
| Fs | 4.48 |  |  | 1.26 |  |  | 0.94 |  |  | 0.55 |  |  | 3.59 |  |  |
| p-value | 0.05 |  |  | 0.30 |  |  | 0.37 |  |  | 0.58 |  |  | 0.02 |  |  |

